# Stress-Induced Dinoflagellate Bioluminescence at the Single Cell Level

**DOI:** 10.1101/2020.03.18.997544

**Authors:** Maziyar Jalaal, Nico Schramma, Antoine Dode, Hélène de Maleprade, Christophe Raufaste, Raymond E. Goldstein

**Affiliations:** Department of Applied Mathematics and Theoretical Physics, University of Cambridge, Cambridge CB3 0WA, United Kingdom; Max-Planck Institute for Dynamics and Self-Organization, Göttingen, Germany; École Polytechnique, 91128 Palaiseau Cedex, France; Université Côte d’Azur, CNRS, Institut de Physique de Nice, CNRS, 06100 Nice, France

## Abstract

One of the characteristic features of many marine dinoflagellates is their bioluminescence, which lights up nighttime breaking waves or seawater sliced by a ship’s prow. While the internal biochemistry of light production by these microorganisms is well established, the manner by which fluid shear or mechanical forces trigger bioluminescence is still poorly understood. We report controlled measurements of the relation between mechanical stress and light production at the single-cell level, using high-speed imaging of micropipette-held cells of the marine dinoflagellate *Pyrocystis lunula* subjected to localized fluid flows or direct indentation. We find a viscoelastic response in which light intensity depends on both the amplitude and rate of deformation, consistent with the action of stretch-activated ion channels. A phenomenological model captures the experimental observations.

---

Bioluminescence, the emission of light by living organisms, has been a source of commentary since ancient times [1], from Aristotle and Pliny the Elder, to Shakespeare and Darwin [2], who, like countless mariners before him, observed of the sea, *“… every part of the surface, which during the day is seen as foam, now glowed with a pale light. The vessel drove before her bows two billows of liquid phosphorus, and in her wake she was followed by a milky train. As far as the eye reached, the crest of every wave was bright,…”*. The glow Darwin observed arose most likely from bacteria or dinoflagellates, unicellular eukaryotes found worldwide in marine and freshwater environments.

Bioluminescence is found in a large range of organisms, from fish to jellyfish, worms, fungi, and fireflies. While discussion continues regarding the ecological significance of light production [3], the *internal* biochemical process that produces light is now well understood. In the particular case of dinoflagellates [4], changes in intracellular calcium levels produce an action potential, opening voltage-gated proton channels in the membranes of organelles called *scintillons*, lowering the pH within them [5] and causing oxidation of the protein *luciferin*, catalyzed by *luciferase*. Far less clear is the mechanism by which fluid motion triggers bioluminescence.

Early experiments on light emission utilizing unquantified fluid stirring or bubbling [6] were superseded over the past two decades by studies in the concentric cylinder geometry of Couette flow [7, 8] and macroscopic contracting flows [9, 10]. Subsequent experiments explored light production by cells carried by fluid flow against barriers in microfluidic chambers [11], or subjected to the localized forces of an atomic force microscope [12]. From these have come estimates of the stress needed to trigger light production. Indeed, dinoflagellates can serve as probes of shear in fluid flows [7, 9, 13–16]. At the molecular level, biochemical interventions have suggested a role for stretch-activated ion channels [17] —known to feature prominently in touch sensation [18] —leading to the hypothesis that fluid motion stretches cellular membranes, forcing channels open and starting the biochemical cascade that produces light.

Here, as a first step toward an in-depth test of this mechanism, we study luminescence of single cells of the dinoflagellate *Pyrocystis lunula* (Fig. 1) induced by precise mechanical stimulation. The cellular response is found to be ‘viscoelastic’, in that it depends not only on the amplitude of cell wall deformation but also on its rate. A phenomenological model linking this behavior to light production provides a quantitative account of these observations.

**FIG. 1.**
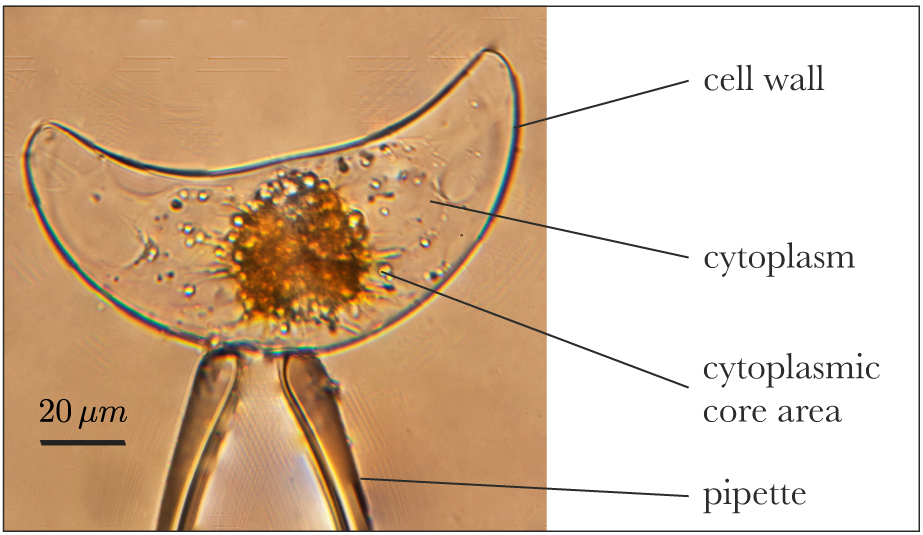
The unicellular marine dinoflagellate *Pyrocystis lunula*, held on a glass micropipette. Chloroplasts (yellow/brown) are in the cytoplasmic core at night and the crescent-shaped cell wall encloses the cell.

*P. lunula* is an excellent organism for the study of bioluminescence because its large size (∼ 40 *µ*m in diameter and ∼ 130 *µ*m in length), lack of motility as an adult, rigid external cell wall and negative buoyancy all facilitate micromanipulation. Its relative transparency and featureless surface allow for high-resolution imaging. As model organisms, dinoflagellates have been studied from a variety of complementary perspectives [19].

Cultures of *P. lunula* (Schütt) obtained from CCAP [20] were grown in L1 medium [21] at 20°C in an incubator on a 12h/12h light/dark cycle. The bioluminescence of *P. lunula* is under circadian regulation [22, 23] and occurs only during the night. All experiments were performed between hours 3 − 5 of the nocturnal phase. An sCMOS camera (Prime 95B, Photometrics) imaged cells through a Nikon 63× water-immersion objective on a Nikon TE2000 inverted microscope. Cells were kept in a 500 *µ*L chamber that allows access by two antiparallel micropipettes held on multi-axis micromanipulators (Patchstar, Scientifica, UK) (see Supplemental Material [24]), and kept undisturbed for several hours prior to stimulation. Upon aspiration on the first pipette, cells typically flash once [25]. Care was taken to achieve consistent positioning of cells for uniformity of light measurements (Video 1 [24]).

The second pipette applies mechanical stimulation in either of two protocols. The first directs a submerged jet of growth medium at the cell, controlled by a syringe pump (PHD2000, Harvard Apparatus) and characterized using Particle Image Velocimetry (PIV) and particle tracking, as in Figs. 2a-f. Typical flow rates through the micropipette were on the order of 1 ml/h, exiting a tip of radius ∼10 *µ*m, yielding maximum jet speeds *U* up to 1 m/s. With *ν* = *η/ρ* = 1 mm^2^/s the kinematic viscosity of water and *𝓁* ∼ 0.02 mm the lateral size of the organism, the Reynolds number is *Re* = *U𝓁/ν* ∼ 20, consistent with prior studies in macroscopic flows [7, 9, 10], which utilized the apparatus scale (mm) for reference. In the second protocol, a cell is held between the two pipettes, and mechanical deformation is imposed by displacement of the second. Using the micromanipulators and a computer-controlled translation stage (DDS220/M, Thorlabs), the deformation *δ* and deformation rate *δ* could be independently varied (Figs. 2g-l).

**FIG. 2.**
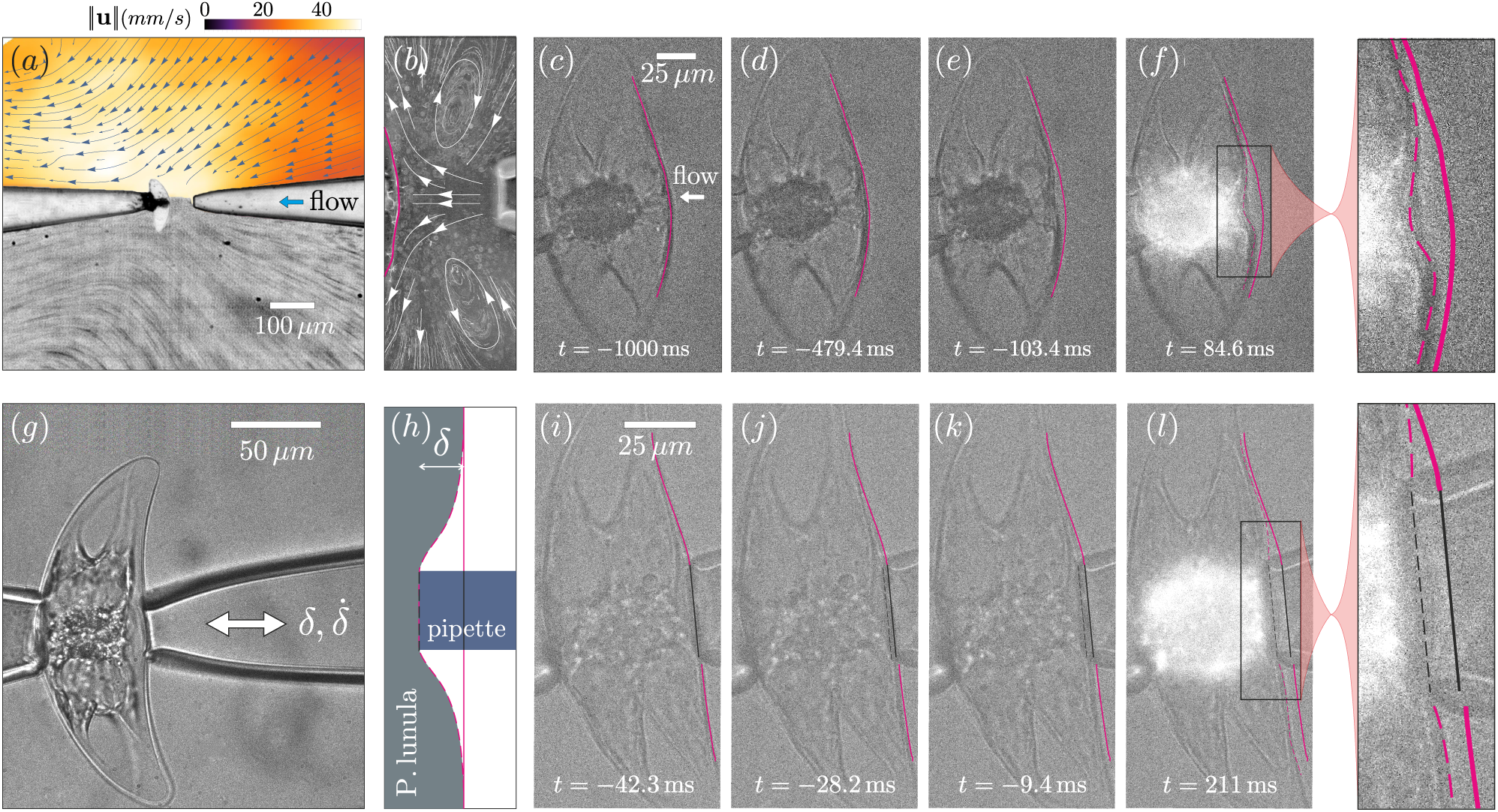
Light production by *P. Lunula* under fluid and mechanical stimulation. (a) Stimulation by fluid flow; color map in upper half indicates flow speed, lower half is a streak image of tracer particles. (b) Particle tracking of flow lines near cell surface. (c-f) Cell deformation due to fluid flow and the consequent light production. (g,h) Second protocol, in which a cell is deformed under direct contact by a second pipette. (i-l) Light production triggered by mechanical deformation. All times indicated are with respect to the start of light emission.

A key observation within the first protocol is that cells do not flash unless the imposed fluid pressure is high enough to deform the cell membrane sufficiently (Fig. 2f). For these submerged jet flows, the fluid stress Σ*_f_* ∼ *ρU* ^2^ can be estimated to reach ∼ 10^3^ Pa, which is the same order as in prior macroscopic experiments [7, 9, 10]. It can be seen from Fig. 2f that the lateral scale *ξ* of cell wall deformations is ∼ 30 *µ*m, and we estimate the fluid force exerted at the site of deformation as *F*_*f*_ ∼ Σ*_f_ ξ*^2^ ∼ 0.1 *µ*N. More quantitatively, using PIV of the flow field and finite-element calculations of the flow from a pipette [24] we find from study of 35 cells that the threshold for light production is broadly distributed, with a peak at 0.10 ± 0.02 *µ*N.

It is not clear *a priori* whether the deformations in Figs. 2c-f are resisted by the cell wall alone or also by the cytoplasm. The wall has a tough outer layer above a region of cellulose fibrils [26–28], with a thickness *d* ∼ 200 − 400 nm: AFM studies [12] show a Young’s modulus *E* ∼ 1 MPa. During asexual reproduction, the cellular contents pull away from the wall and eventually exit it through a hole, leaving behind a rigid shell with the characteristic crescent moon shape [29]. Thus, the wall is not only imprinted with that shape, but is much more rigid than the plasma membrane and significantly more rigid than the cytoplasm [12].

Deformations of such curved structures induced by localized forces involve bending and stretching of the wall. With *𝓁* the radius of curvature of the undeformed cell wall, a standard analysis [30] gives the indentation force *F* ∼ *Ed*^2^*δ/𝓁*. Balancing this against the fluid force *ρU*^2^ *ξ*^2^ we find the strain *ε* ≡ *δ/𝓁* ∼ (*ρU*^2^ */E)* (*ξ/d*)^2^. From the estimates above, we have *ρU* ^2^*/E* ∼ 10^−4^, and *ξ/d* ∼ 50 − 100, so *ε* is of the magnitude observed.

In the natural setting of marine bioluminescence and in laboratory studies of dilute suspensions, light production can arise purely from flow itself, without contact between dinoflagellates. Nevertheless, there are conceptual and methodological advantages to studying bioluminescence by direct mechanical contact, especially due to the natural compliance of cells aspirated by a single micropipette. Chief among these is the ability to control the deformation and deformation rate, which are the most natural variables for quantification of membrane stretching and bending. As seen in Fig. 2i-l, imposing deformations similar to those achieved with the fluid flow also produces bioluminescence, highlighting the role of cell membrane deformation in mechanosensing.

In our protocol for deformations, *δ* is increased at a constant rate *δ*. for a time *t*_*f*_ to a final value *δ*_*f*_ (*loading*), after which it was held fixed until any light production ceases, then returned to zero (*unloading*). We observe generally that if light is produced during loading, it is also produced during unloading. Experiments were performed for *δ*_*f*_ ∈ [1, 10] *µ*m and 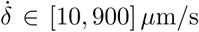, with eight to twelve replicates (cells) for each data point. We repeated the given deformation protocol on the same cell (with sufficient rest intervals in between) until the cell ceases bioluminescence. Reported values of light intensity *I*(*t*) are those integrated over the entire cell.

Figure 3a shows the light flashes from 15 stimulations of a single cell. With each deformation, *I*(*t*) first rises rapidly and then decays on a longer time scale. Apart from a decreasing overall magnitude with successive flashes, the shape of the signal remains nearly constant. The eventual loss of bioluminescence most likely arises from exhaustion of the luciferin pool [31]. The inset shows the corresponding phase portraits of the flashes in the *I* − *dI/dt* plane, where the similarity of successive signals can clearly be seen.

**FIG. 3.**
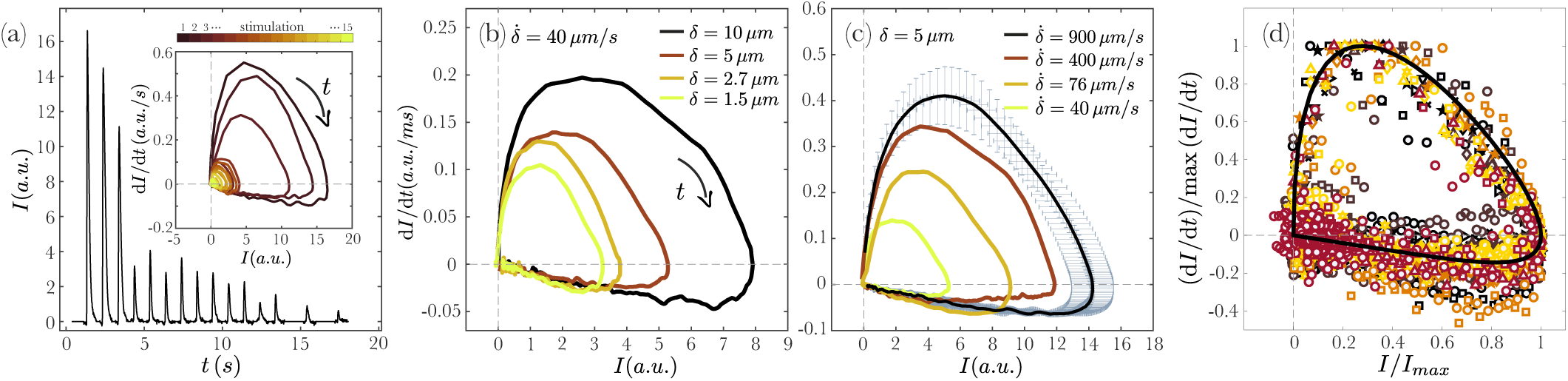
Dynamics of light production following mechanical stimulation. (a) Response of a cell to repeated deformation with *δ*_*f*_ = 10 *µ*m and 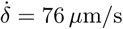. Inset: loops in *I* − *dI/dt* plane for successive flashes. (b) Loops at fixed 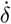 and varying *δ*_*f*_ for first flashes. (c) As in (b), but for fixed *δ*_*f*_ and varying 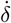. Standard errors are shown for outermost data. (d) Master plot of data, normalized by maximum intensities and rates. Circles (squares) are data in b (c). Black curve is result of model in (1) and (3).

Focusing on the first flashes, experiments with different *δ*_*f*_ and 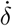 reveal the systematics of light production. Figures 3b&c show that for a given rate, larger deformations produce more light, as do higher rates at a given deformation. Interestingly, the shape of the signals remains the same not only between different cells but also for different mechanical stimulations; normalizing the phase portraits with respect to their maxima yields a universal shape of the signal (Fig. 3d). We summarize the results of all experiments in Figure 4a, showing the variation of maximum light intensity (averaged over all the first flashes) as a function of *δ*_*f*_ and 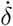; light production is maximized when the cell is highly deformed at high speed.

**FIG. 4.**
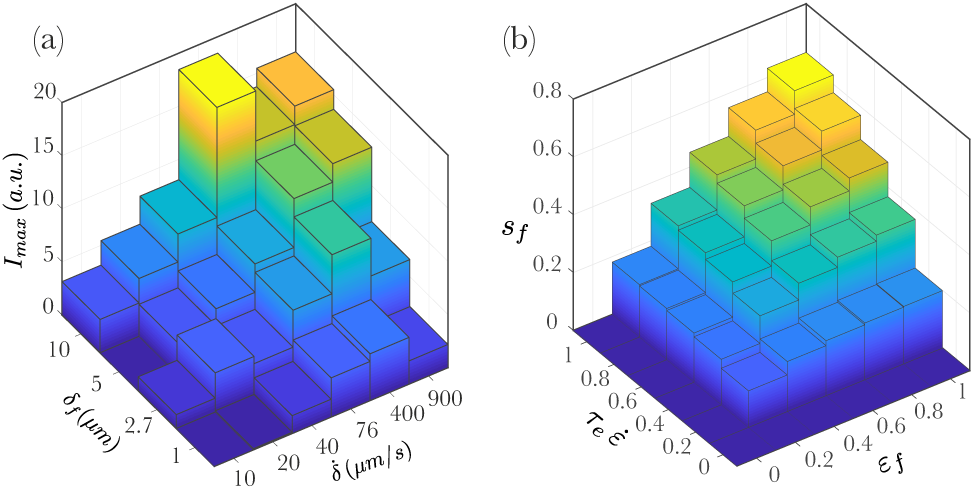
Dependence of light production on deformation and rate. (a) Histogram of maximum intensity. Note nonuniform grid. (b) Variation of signal strength *s*_*f*_ predicted by phenomenological model, as a function of deformation and rate.

The influence of deformation and rate are suggestive of viscoelastic properties. At a phenomenological level, we thus consider a Maxwell-like model that relates the signal *s*(*t*) that triggers light production to the strain *ε*,

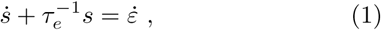

where *τ*_*e*_ is a relaxation time. For a given *δ*, if the deformation time scale is much smaller than *τ*_*e*_, the membrane does not have time to re-arrange (the large Deborah number regime in rheology), while for slow deformations the membrane has time to relax. As seen in Figs. 2i-l and Videos 2 & 3 [24], bioluminescence occurs *during* loading, a feature that suggests *τ*_*e*_ is comparable to the flash rise time. Integrating (1) up to *t*_*f*_, we obtain the signal *s*_*f*_ at the end of loading in terms of the final strain *ε*_*f*_ ≡ *δ*_*f*_ /*l* and scaled strain rate 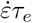,

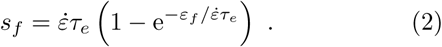

As seen in Fig. 4b, the peak response occurs when both the final strain and strain rate are large, as observed experimentally. The linear relationship between *s* and *ε* embodied in (1) can not continue to be valid at large strains or strain rates; eventually, the signal must saturate when all available channels open to their maximum. This is consistent with the data in Fig. 4a at the highest rates, where experimentally *ε* ∼ 0.25.

Although light production is triggered internally by an action potential—which arises from nonlinear, *excitable* dynamics—analysis of the flashes [24] indicates a time course much like that of two coupled capacitors charging and discharging on different time scales. Such linear dynamics have figured in a variety of contexts, including calcium oscillations [32], bacterial chemotaxis [33], and algal phototaxis [34], and take the form of coupled equations for the observable (here, the light intensity *I*) which reacts to the signal *s* on a short time *τ*_*r*_ and the hidden biochemical process *h* which resets the system on a longer time *τ*_*a*_. For light triggered by stretch-activated ion channels, the signal *s* might be the influx of calcium resulting from the opening of channels. Adopting units in which *I, h, s* are dimensionless, the simplest model is

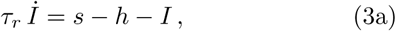

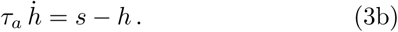

Starting from the fixed point (*I* = 0, *h* = 0) for *s* = 0, if the signal is turned on abruptly then *I* will respond on a time scale *τ*_*r*_, exponentially approaching *s* − *h* ≃ *s*. Then, as *h* evolves toward *s* over the longer adaptation time scale *τ*_*a*_, *I* will relax toward *s* − *h* ≃ 0, completing a flash. It follows from (3) that a discontinuous initial *s* creates a discontinuous *İ*, whereas the loops in Fig. 3 show smooth behaviour in that early regime (*I, İ≳* 0); this smoothing arises directly from the Maxwell model for the signal. The parsimony of the linear model (3) comes at a cost, for it fails at very high ramp rates when 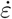 switches to zero within the flash period and both *s* and *I* would adjust accordingly, contrary to observations. In a more complex, excitable model, the flash, once triggered, would thereafter be insensitive to the signal.

As the entire system (1) and (3) is linear, it can be solved exactly [24], thus enabling a global fit to the parameters. We compare the theoretical results with the normalized experimental data in Fig. 3d, where we see good agreement with the common loop structure. From the fits across all data, we find common time scales *τ*_*e*_ ≈ 0.027 s, *τ*_*r*_ ≈ 0.012 s, and *τ*_*a*_ ≈ 0.14 s, the last of which is comparable to the pulse decay time found in earlier experiments with mechanical stimulation [25], and can be read off directly from the late-time dynamics of the loops in Figs. 3b&c, where *İ* ∼ −*I/τ_a_* [24]. These values suggest comparable time scales of membrane/channel viscoelasticity and biochemical actuation, both much shorter than the decay of light flashes.

With the results described here, the generation of bioluminescence has now been explored with techniques ranging from atomic force microscope cantilevers with attached microspheres indenting cells in highly localized areas, to fluid jets and micropipette indentation on intermediate length scales, and finally to macroscopic flows that produce shear stresses across the entire cell body. Figure 5 considers all of these experiments together, organized by the perturbative stress Σ found necessary to produce light and the area A ≡ *ξ*^2^ over which that stress was applied. We see a clear trend; the smaller the perturbation area, the larger the force required. This result suggests that the production of a given amount of light, through the triggering effects of stretch-activated ion channels on intracellular action potentials, can be achieved through the action of many channels weakly activated or a small number strongly activated. With an eye toward connecting the present results to the familiar marine context of light production, it is thus of interest to understand more quantitatively the distribution of forces over the entire cell body in strong shear flows [35] and how those forces activate ion channels to produce light. Likewise, the possible ecological significance of the great range of possible excitation scales illustrated in Fig. 5 remains to be explored.

**FIG. 5.**
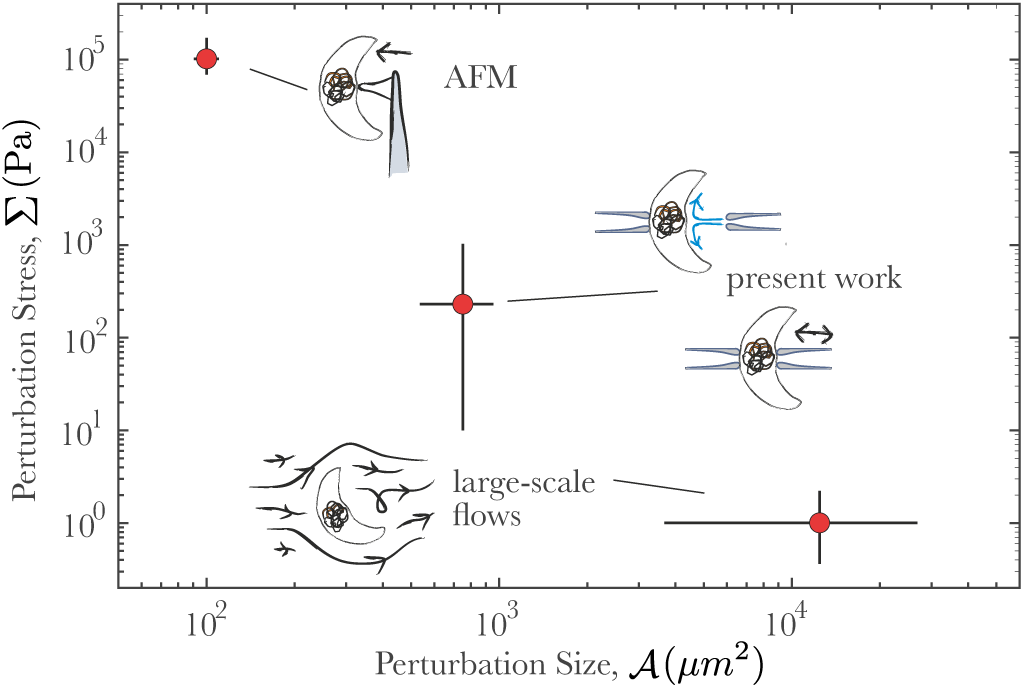
Perturbation stress versus perturbation area for three kinds of experiments on dinoflagellates. Atomic force measurements on *P. lunula* are from [12], while macroscopic measurements include Taylor-Couette [7, 8] and contracting flows [9, 10] on *P. lunula* and similar dinoflagellates.

We are grateful to Michael I. Latz for invaluable assistance at an early stage of this work, particularly with regard to culturing dinoflagellates, and thank Adrian Bar-brook, Martin Chalfie, Michael Gomez, Tulle Hazelrigg, Chris Howe, Caroline Kemp, Eric Lauga, Benjamin Mau-roy, Carola Seyfert, and Albane Théry for important discussions. This work was supported in part by the Gordon and Betty Moore Foundation (Grant 7523) and the Schlumberger Chair Fund. CR acknowledges support by the French government, through the UCAJEDI Investments in the Future project of the National Research Agency (ANR) (ANR-15-IDEX-01).

