## Supplemental Material for "Stress-Induced Dinoflagellate Bioluminescence at the Single Cell Level"

### I. EXPERIMENTAL SETUP

The experimental setup, shown schematically in Figure S1, consists of a microscope for visualization and positioning systems to control the two pipettes. All experiments were conducted in a darkened room. The white light illumination of the microscope (Nikon TE2000 ) was kept to a minimum and sent through a red long-pass filter (620 nm, Knight Optical, UK) to avoid disruption of the night phase of the dinoflagellates and to allow a greater dynamic range in capturing the bioluminescence. That background intensity was controlled in all experiments for uniformity.

Pipettes were positioned with 3D micromanipulators (Patchstar, Scientifica). For small deformation rate experiments, we used a Thorlabs 1D Direct Drive Linear Stage (DDS220/M) to control the motion. All stages were programmed with their native software. The pipettes were connected to syringes with stiff tubing and fluid flow through them could also be used to position the cells (see below). In the flow experiments, we used a syringe pump (PHD2000, Harvard Apparatus) at a constant rate. The test section was a chamber whose top and bottom were coverslips, held apart by $\sim 2$ mm plastic spacers (see inset of Fig. S1). As *P. lunula* has a very characteristic three dimensional geometry, for consistency, we held the cells the same orientation within the chamber in all experiments.

As cells of *P. lunula* are negatively buoyant, they settle to the bottom coverslip of the sample chamber. Cells were positioned manually with the use of joystick controllers and gentle suction of the flow, as illustrated in Figure S2. The main bioluminescence experiments were recorded with a Prime 95B sCMOS camera (Photometrics). The high sensitivity of the camera allowed for measurements at low light condition but relatively high recording speed. For the PIV and particle tracking experiments, we used a high-speed camera (Phantom v311). Figure 1 of the main text was

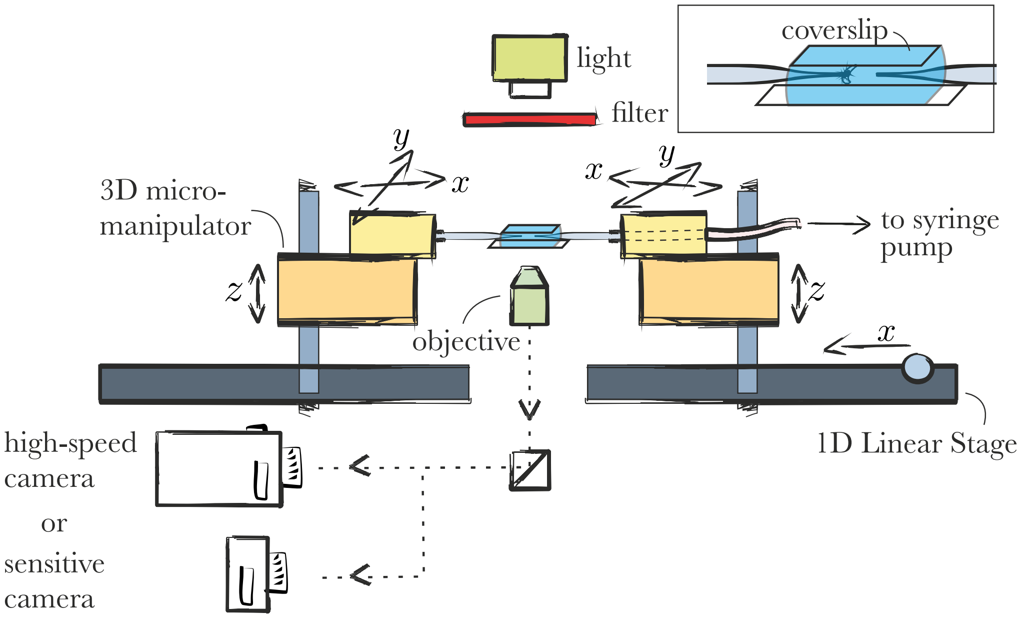


FIG. S1: Schematic of experimental setup to study bioluminescence produced by single dinoflagellates under controlled stresses.

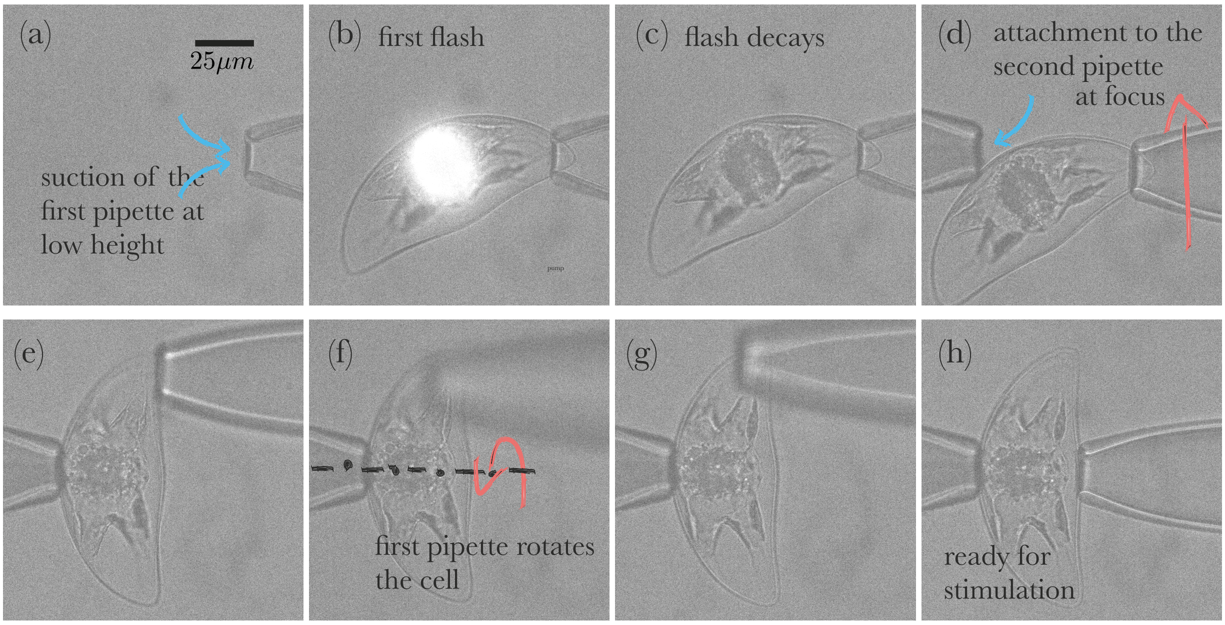


FIG. S2: Manual positioning of a cell prior to main measurements. (a) The cell is initially drawn up from the bottom cover glass using gentle flow suction. It nearly always aspirated from one of its pointy ends. The cell flashes once in this process (b) and the light decays (c). The pipettes are raised within the sample chamber to be far from the top and bottom chamber surfaces. (d) Using the joystick controllers of the micromanipulators, the cell is placed on the other pipette and then held using gentle suction (e). (f,g) The cell is rotated so that the largest area is exposed to the camera. (h) Finally, by placing the cell between the pipettes, indentation experiments can be performed.

captured using a Nikon D810 DSLR with Differential Interference Contrast (DIC) microscopy.

### II. SOLUTION OF THE MODEL

Equations 1, 3a and 3b are linear ODEs which can be solved exactly. As described in the main text, we take here the simplest case in which the light flash occurs within the ramp period, and therefore confine the discussion to times $t < t_f$, during which the rate of strain $\dot{\varepsilon}$ is constant. From the three time constants $(\tau_r, \tau_e, \tau_a)$ we find $\tau_a$ to be by far the largest, and thus define the two ratios $\lambda, \rho < 1$,

$$\lambda = \frac{\tau_e}{\tau_a}\,, \qquad \rho = \frac{\tau_r}{\tau_a}\,. \tag{S1}$$

Then, from (1) the signal is

$$s(t) = \dot{\varepsilon}\tau_e \left(1 - e^{-t/\tau_e}\right)\,. \tag{S2}$$

As $t \to 0$, $s \sim \dot{\varepsilon}t + \cdots$, while at long times $s$ approaches $\dot{\varepsilon}\tau_e$. If we set $t = t_f$ and note that $t_f \equiv \varepsilon_f/\dot{\varepsilon}$, we obtain (2) in the main text. Substituting (S2) into (3b) and solving for $h$, we find

$$h = \frac{\dot{\varepsilon}\tau_e}{1-\lambda}\left[1 - e^{-t/\tau_a} - \lambda\left(1 - e^{-t/\tau_e}\right)\right]\,. \tag{S3}$$

which varies as $\dot{\varepsilon}t^2/2\,\tau_a$ as $t \to 0$ and, as with $s$, approaches $\dot{\varepsilon}\tau_e$ for long times.

Finally, the light intensity is

$$I(t) = \frac{\dot{\varepsilon}\tau_e}{1-\lambda}\left[\frac{1}{1-\rho}\left(e^{-t/\tau_a} - e^{-t/\tau_r}\right) - \frac{\lambda}{\lambda-\rho}\left(e^{-t/\tau_e} - e^{-t/\tau_r}\right)\right], \tag{S4}$$

which behaves as $I \sim \dot{\varepsilon}t^2/2\,\tau_r$ as $t \to 0$. At large times, with $\tau_a > \tau_e \sim \tau_r$, the dominant term in (S4) is $I \propto e^{-t/\tau_a}$, so $\dot{I} \sim -I/\tau_a$, a relationship seen in Figs. 3b&c of the main text. Figure S3 shows plots of the solutions above.

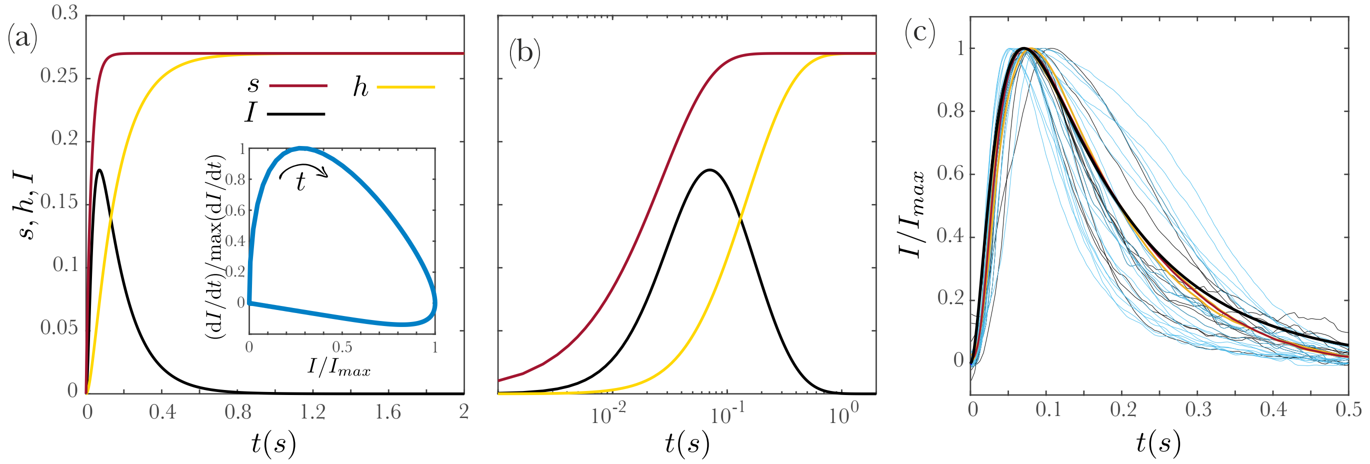


FIG. S3: a) Plots of $s, h$, and $I$ for $\tau_r = 0.012$, $\tau_a = 0.14$, $\tau_e = 0.027$, and $\dot{\delta} = 500\mu$m/s. The inset shows the phase portrait of the intensity signal. b) The same as (a) in a linear-log scale to highlight the early time behaviour. c) Intensity signals corresponding to the results shown in figure 3. Black and blue thin lines show the raw data for fixed $\dot{\delta}$ (3a) and $\delta$ (3b) and $\dot{\delta}$ (3a), respectively, and the yellow lines show their average values. The red line shows the average value of all the raw data. The black line is the same theoretical curve shown in panels a and b, normalized by its maximum value.

To find the time scales $\tau_r, \tau_e$ and $\tau_a$, we employed a least squares analysis on the average signal from all experiments. The values obtained, $\tau_e \approx 0.027$ s, $\tau_r \approx 0.012$ s, and $\tau_a \approx 0.14$ s, yield the ratios $\lambda \approx 0.19$ and $\rho \approx 0.09$. Thus, the prefactors within square brackets in (S4) are $1/(1-\rho) \approx 1.1$ and $\lambda/(\lambda-\rho) \approx 1.9$. Figure S3c compares the theoretical curve for the flash intensity with the experimental data used in Figure 3 of the main text.

### III. FINITE ELEMENT COMPUTATIONS

We performed experiments and counterpart numerical computations for 35 cells to estimate the force required for light production. The steady-state axisymmetric Navier-Stokes equations were solved numerically with the finite element software COMSOL [S1] to obtain the flow from a pipette impinging on a cell. Figure S4a shows a close up of the geometry employed. The geometry of the dinoflagellate is simplified to a sphere of radius $\mathcal{R}$, positioned a distance $H$ from the outlet of the pipette. The computational domain was chosen to be sufficiently large that the presence of the domain boundaries did not affect the calculations. The inner diameter of the micropipette nozzle was set at $25\,\mu$m, with a flow rate $Q = 1$ ml/h, resulting in a fluid exit speed from the micropipette of $V \sim 1$ m/s. Based on the actual size of the organisms, and its distance from the pipette, we performed the simulations for an average value of $\mathcal{R} = 30\,\mu$m. The computations were found to be insensitive to changes in $\mathcal{R}$ within our experimental values. The values of $H$ varied between 9 and 75 $\mu$m.

We compared PIV measurements of the flow created by the submerged jet in the absence of the dinoflagellate to the flow field computed within COMSOL, and found a good agreement. After this validation, we computed the fluid flow in the presence of the sphere (see Fig. S4) and determined the mechanical forces exerted on the surface of the dinoflagellate (sphere) by integrating the stress over its surface. By synthesizing these results with the experimental thresholds for light production we obtain in Figure S4b the probability distribution of the threshold force for bioluminescent flashes. The distribution peaks at $F_f \sim 0.1\,\mu$N. This value is consistent with the estimation based on the dynamic pressure $\Sigma_f$ presented in the main text.

[S1] *COMSOL, Multiphysics v. 5.3., COMSOL AB, Stockholm, Sweden.*

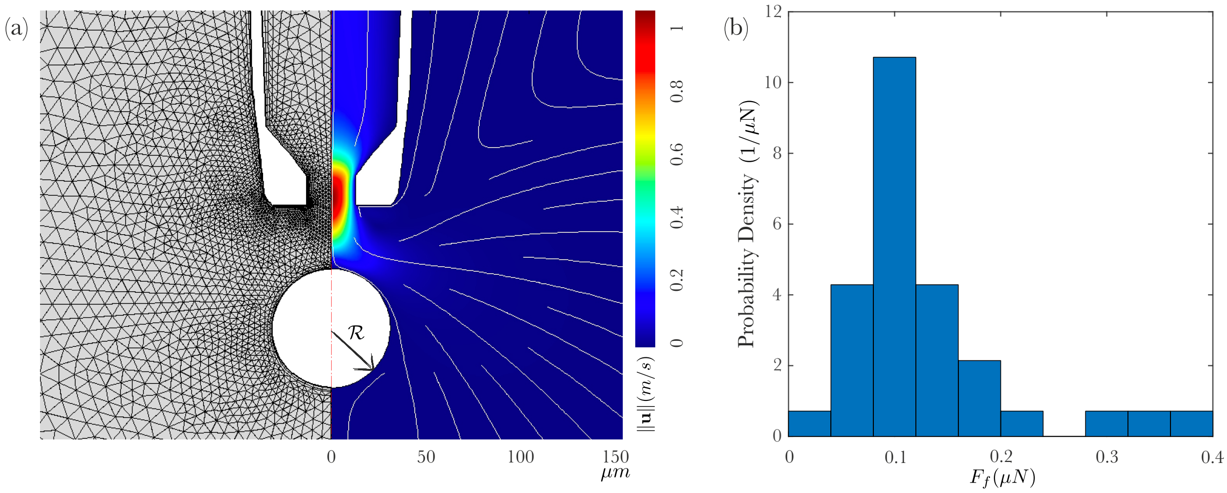


FIG. S4: (a) Numerical simulation of the fluid flow around an organism, modeled as a sphere: (left) mesh, (right) velocity magnitude. (b) Histogram of the threshold force for light production.
